# Snaq: A Dynamic Snakemake Pipeline for Microbiome data analysis with QIIME2

**DOI:** 10.1101/2022.03.10.483866

**Authors:** Attayeb Mohsen, Yi-An Chen, Rodolfo S. Allendes Osorio, Chihiro Higuchi, Kenji Mizuguchi

## Abstract

Optimizing a protocol for 16S microbiome data analysis with QIIME2 is a challenging task for biologists with no programming experience. It involves a multi-step process, and multiple parameters and options that need to be tested and determined. In this article, we describe Snaq, a snakemake pipeline that helps automate and optimize 16S data analysis using QIIME2. Snaq offers an informative file naming system and automatically performs the analysis of a data set by downloading and installing the required databases and classifiers, all through a single command-line instruction. It works natively on Linux and Mac and on Windows through the use of containers, and is potentially extendable by adding new rules. This pipeline will substantially reduce the efforts in sending commands and prevent the confusion caused by the accumulation of analysis results due to testing multiple parameters.

## Introduction

The microbial content of a biological sample can be determined by sequencing and the subsequent bioinformatic processing/analysis of the sequenced data. 16S Ribosomal RNA gene sequencing (Hugerth and Andersson 2017) is one of the most intensively used approaches in microbiome research. It is also called amplicon sequencing since it incorporates the amplification of a specific DNA region (16S RNA gene) in bacterial genomes using PCR. There are multiple bioinformatics tools available for the analysis of this type of data (Gołębiewski and Tretyn 2019; Prodan et al. 2020).

QIIME2 (Bolyen et al. 2019) is a microbiome data analysis platform that targets amplicon (16S) data. By allowing the integration of different software programs, implemented as plugins (such as DADA2 for data denoising), QIIME2 is designed to facilitate seamless incorporation of new plugins, allowing developers to add new features easily^1^.

QIIME2 plugins handle input and output through the definition of *artifacts*, i.e., compressed folders that contain both data files and metadata information. For example, raw sequence data can be imported to construct an artifact, which is later used by a specific plugin. In turn, the plugin produces a new artifact of a different type as output.

This approach makes it possible to change the order of the steps or insert new steps in the middle without extra effort, provided that the input and output follow the QIIME2 framework guidelines. This approach makes combining multiple tools in sequence effortless and reduces the requirements of programming skills. Moreover, importing or exporting data from/to various formats or visualization of the original data or the results can be achieved easily.

Despite its multiple advantages, some difficulties arise when trying to automate data analysis using QIIME2. Even when the same data types and the same experimental technique are used, the analysis results depend on multiple environmental and technical features, such as the length and quality of the sequenced data. As the choice of the tools and parameters used for the analysis depend on the data set, every data set can be considered unique and in need of special treatment, making it impossible (or very difficult) to automate.

For example, the process of *quality trimming* is used to clean the data and improve the results by removing the nucleotides assigned with low confidence. Selecting the appropriate quality threshold value at which trimming should be performed depends on the data set, and usually requires trying and testing. If a very stringent trimming threshold is adopted, plenty of good data could be lost; alternatively, adopting a loose trimming threshold introduces low-quality data in the downstream analysis, affecting the quality and reliability of the results. Moreover, there are different tools and databases for data cleaning, identifying Operational Taxonomy Units (OTUs), and taxonomy assignment.

Usually, researchers need to investigate multiple options, which requires running the analysis several times and comparing the results to decide the best set of tools and parameters for the data under investigation. Such an optimization process requires a substantial effort and leads to the accumulation of several copies of the results with different sets of parameters, making the whole process rather inefficient and difficult to reproduce.

Although some pipelines for the automated analysis of 16S data are available (Estaki et al. 2020; Fung et al. 2021; Kondratenko et al. 2020; Hu and Alexander 2020), none of them provide an easy way to run the analysis multiple times for optimization purposes. Also, they tend to have fixed steps and do not make it easier to change the sequence of steps and parameters used.

Snakemake (Koster and Rahmann 2012; Mölder et al. 2021) is a Python dialect created for the specification of pipeline workflows. A Snakemake pipeline is specified through the definition of *rules*; where each rule typically has: *input* for the specification of input files; *output* for the specification of output files; and *shell* for the specification of the command used to produce the output based on the input.

The execution of a Snakemake pipeline is achieved via the definition of a single target file name. Snakemake will then determine the steps required to produce the target output based on its rules, the file name, and the application of the wildcards concept^2^.

The wildcards concept facilitates passing parameters for any rule in the pipeline by inferring the parameter’s value from the target file name. This feature of Snakemake is especially suitable for parameter optimization. There are other advanced features, such as caching processing results, to prevent doing the same analysis repeatedly (Koster and Rahmann 2012).

By combining the analysis strengths of QIIME2 with the flexibility in the definition of pipelines provided by Snakemake, here we introduce “Snaq”, a dynamic Snakemake pipeline for microbiome data analysis with QIIME2.

Snaq incorporates the definition of analysis rules with the definition of an expressive target file format, which together provide the functionality required to achieve the following when working with QIIME2:

1. **Easy protocol optimization**: By changing the name of the target file, the analysis workflow dynamically changes, allowing testing of different tools and parameters. This is crucial, as the analysis of 16S microbiome data with QIIME2 can be performed in multiple ways with numerous permutations of software and parameter choices, depending on the technology used and sequencing qualities, and the researcher’s preference.
2. **Full pipeline automation:** Combining rule definition with an ad-hoc target file name, Snaq allows the execution of a full analysis pipeline through a single command instruction. As 16S microbiome data analysis with QIIME2 entails multiple command submission, this significantly reduces the number of commands and instructions that the user needs to know, allowing to focus on the actual analysis and not the programming.
3. **Handle data accumulation:** Snaq automatically handles the (intermediate) data that are often generated as a result of multiple trial runs. Additionally, it avoids the duplication of intermediate result files when multiple executions of different analysis pipelines include identical intermediate stages.

## Implementation

Snaq is made of three main components: 1) `Snakefile`, 2) `env` folder and 3) `script` folder. `Snakefile` is the file where all the required snakemake rules are implemented; notice these rules were carefully constructed as not to contend with each other and to make the whole process run smoothly. The `env` folder contains the definitions for the Conda environments as a series of YAML files, while the `scripts` folder contains extra scripts required by Snaq to fill the gaps of the pipeline that are not covered by QIIME2 plugins.

Snaq takes advantage of QIIME2’s command-line interface and available plugins and combines it with our implementation of new Python and R scrips. Then, by incorporating a descriptive name file convention and the rule-based structure of Snakemake, it makes possible the definition and execution of dynamically defined pipelines through a single terminal command.

Snaq can be used on personal computers or server environments. It works on Linux and Mac operating systems^3^. It is also possible to use directly from the available Docker and singularity containers. All analysis takes place in a Snaq folder, where the input files need to be stored inside `data/` folder, and all results will be saved in `results/` folder.

### Descriptive File Convention

To make the pipeline versatile and easily modifiable, we adopted a convention of including all the key parameter values inside a target file name and called this scheme descriptive target file naming (Figure: 1). This means that Snakemake will parse the target file name and infer the sequence of steps and the parameter values used. Then the target file will be created accordingly.

**Figure 1:**
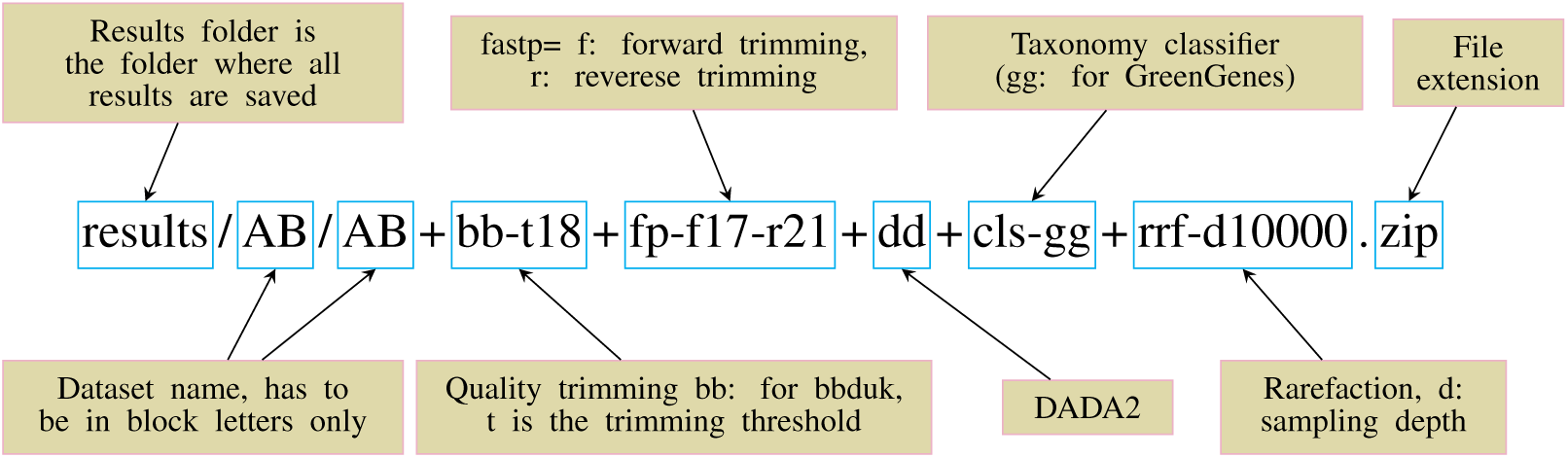
Descriptive target file name example. A target file is required for Snaq to execute correctly.

To let Snakemake infer the required steps and their order, we used a predetermined output nomenclature for each stage (Table 1).

**Table 1:**
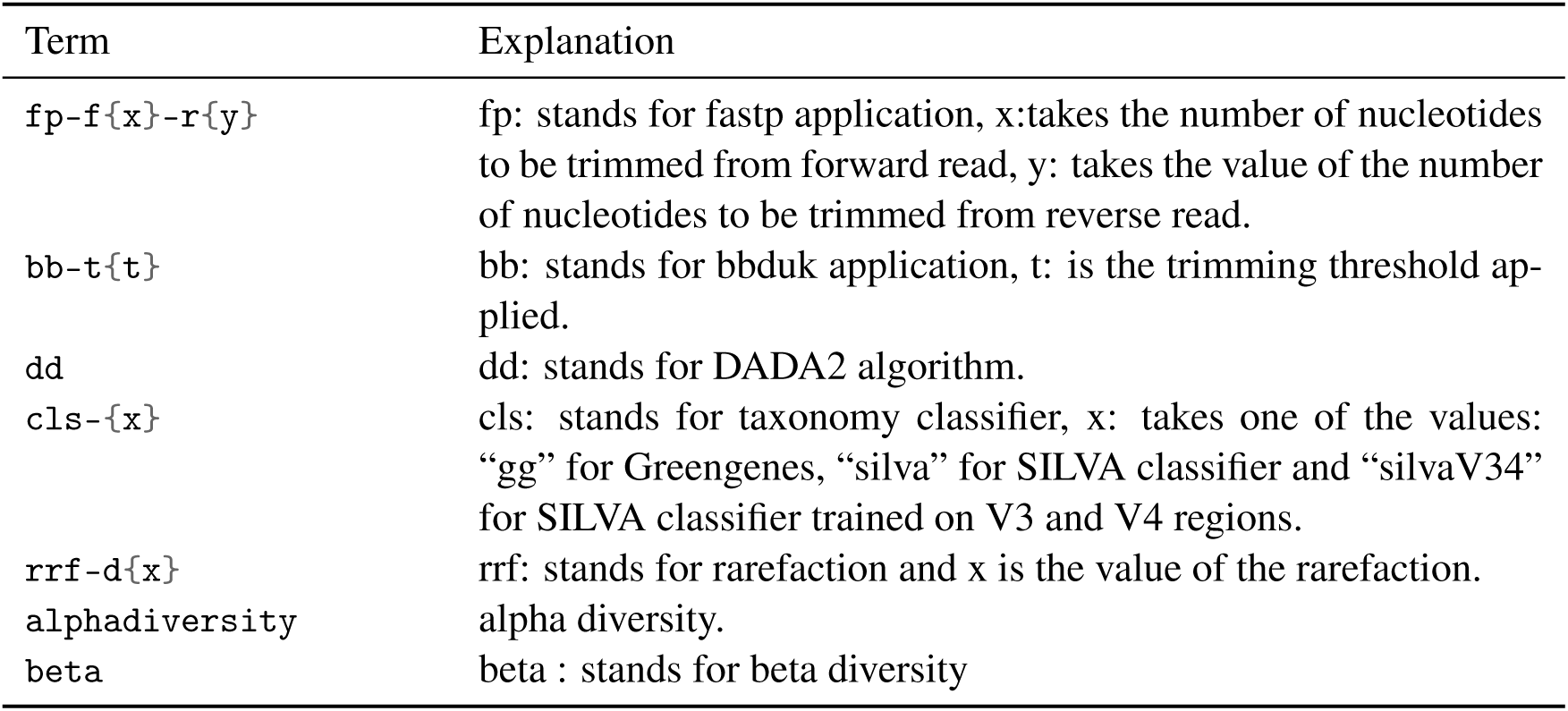
Output Nomenclature

The stages of analysis in the target file name are divided using the character “+” (plus sign). For example, let us consider the case of the target file shown in Figure 1. Here, we are requesting Snaq to produce a summarized result (indicated by the extension `.zip`) for the input data located in folder `data/AB/`^4^. Sequentially, `bb-t18` indicates a trimming stage with threshold value 18; `fp-f17-r21` indicates the use of fastp with a forward trimming value of 17 and a reverse trimming value of 21; `dd` indicates the use of the DADA2 algorithm; `cls-gg` request a taxonomy classification using Greengenes; and finally `rrf-d10000` indicates the use of rarefaction with a sampling depth of 10000.

We believe that in most cases, users will use Snaq through the definition of a single target file and a single command-line instruction. However, intermediate results files can also be produced upon request by using the corresponding target file. For example, if a user were only to import the dataset into a QIIME2 artifact, this could be done by using `results/AB/AB.qza` as target file name. Similarly, if trimming were to be added, the corresponding target file would be `results/AB/AB+bb-t18.qza`, and so on. Notice that the addition of stages is typically in a forward fashion; this means that later analysis stages can not be added to the target file name without their previous stages also being part of it.

The multiple stages specified in the target file name define an execution pipeline, as the one shown in Figure 2.

**Figure 2:**
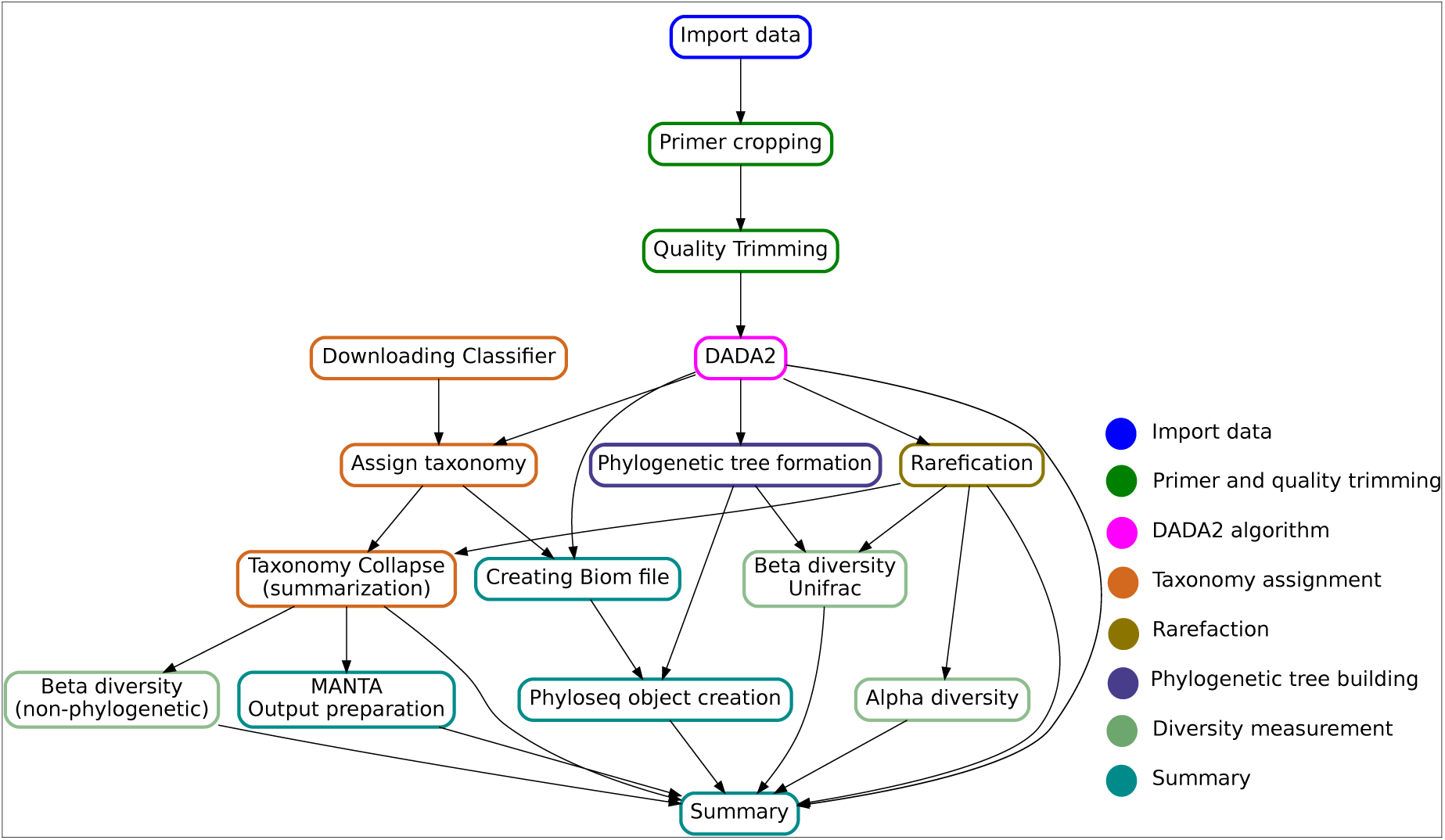
Directed Acyclic Graph showing the key steps in the Snaq pipeline. Notice that the given target file will determine a specific traversal of the graph. Color of nodes is used to indicate the analysis stage, as proposed in the implementation section.

In the following, we provide a detailed description of the pipeline stages and their corresponding^5^:

- **Import data**: This stage imports FASTQ files from the source folder to a QIIME2 artifact (qza). Notice that, to avoid the dataset needs to be named using only capital letters. The results for this step are stored in folder `results/AB/AB.qza`. The command required to run this step is: `snakemake --use-conda --cores 10 results/AB/AB.qza` Hereafter, we will omit the command and options (`snakemake --use-conda --cores 10`), and focus only on the target file name.
- **Primer trimming:** This stage uses fastp (Chen et al. 2018) to trim a specified number of nucleotides in both reads. The format of the target is `fp-f{X}-r{Y}` where `X` represents the number of nucleotides to be cropped from the 5’ end of the forward reads (R1), and `Y` is the number of nucleotides to be cropped from the reverse reads (R2). For example, in order to add the trimming of 17 bases from the 5’ end of R1 and 21 bases from R2 to our previously loaded dataset, the target would be: `results/AB/AB+fp-f17-r21.qza`
- **Quality trimming:** It uses bbduk (part of the bbmap tools) (Bushnell 2021) to trim the section with low quality at the end of the reads in both R1 and R2. The format target for this step is `bb-t{X}` where `X` represents the trimming threshold. To add a quality trimming of reads with threshold of 18, the target file name becomes: `results/AB/AB+fp-f17-r21+bb-t18.qza` Both primer and quality trimming procedures are optional (can be omitted) and their order can be reversed. For example, the following target file names are also valid: `results/AB/AB+bb-t18.qza` `results/AB/AB+bb-t18+fp-f17-r21.qza`
- **DADA2 algorithm:** The DADA2 (Callahan, McMurdie, Rosen, et al. 2016) stage filters the reads, joins pairs, and removes chimera producing Amplicon sequence variant tables (ASVs) that replace OTUs in traditional clustering methods such as UCLUST (Callahan, McMurdie, and Holmes 2017). As result, three different outputs are generated: an Amplicon sequence variant (ASV) frequency table (`dd_table.qza`), a table of representative sequences for ASVs (`dd_seq.qza`) and the statistics of DADA2 performance (`dd_stats.qza`). Using any one of as target will trigger the generation of all three files, for example: `results/AB/AB+fp-f17-r21+bb-t18+dd_seq.qza` When the DADA2 algorithm stage is part of a longer pipeline, its inclusion in the target file name can be simply identified by using the word `dd` (see next section’s target file name).
- **Taxonomy assignment:** It uses the “feature-classifier” plugin (Robeson et al. 2020; Bokulich et al. 2018) to predict the taxonomy of ASVs. Three different classifiers are available: Greengenes (`cls-gg`) (DeSantis et al. 2006; McDonald et al. 2011), SILVA (`cls-silva`) (Pruesse et al. 2007; Glöckner et al. 2017), and SILVA trained on V3 and V4 regions (`cls-silvaV34`) (Mohsen 2021). The resulting output can be generated both as a QIIME2 artifact (`cls-<classifier>_taxonomy.qza`) or as tab separated file (`cls-<classifier>_taxonomy.tsv`). For the Greengenes classifier, the two target file name alternatives would be: `results/AB/AB+fp-f17-r21+bb-t18+dd+cls-gg_taxonomy.qza; and` `results/AB/AB+fp-f17-r21+bb-t18+dd+cls-gg_taxonomy.tsv`
- **Phylogenetic tree building:** This step uses the fasttree algorithm (Price et al. 2010) and QIIME2 phylogeny plugin (Qiime2 Developers 2021) to produce a phylogenetic tree file in `NWK` format (using `fasttree.nwk` as target) or QIIME2 artifact (using `fasttree_rooted.qza` as target). Notice that, since the building of a phylogenetic tree can be done directly after the DADA2 algorithm, the following is a valid target file name: `results/AB/AB+bb-t18+fp-f17-r21+dd+fasttree.nwk`
- **Rarefaction**: The inclusion of a rarefication stage is indicated by using the `rrf-d{X}` target, where `X` represents the sampling depth as defined in (Hughes and Hellmann 2005). Notice that rarefaction needs to be applied before alpha diversity, nonphylogenic beta diversity measurements or the generation of biom tables. To apply rarefaciton the following target file names can be used: `results/AB/AB+bb-t18+fp-f17-r21+dd_table+rrf-d10000.qza`, to generate a QIIME2 artifact, or `results/AB/AB+bb-t18+fp-f17-r21+dd_table+rrf-d10000.tsv`, for a tab separated value file. In this step, the part `_table` was not omitted because rarefaction affects only the table and because in the following stages, this rarefied table is to be used to create biom tables and manta files.
- **Diversity measurement:** At this stage, QIIME2 is used for the computation of alpha and beta diversities. Whilst the target for alpha diversity is simply `alphadiversity`, different types of beta diversity are specified through the target `beta_<type>`, where `<type>` is one of the following: `braycurtis`, `jaccard`, `unweightedunifrac` or `weightedunifrac`. Sample target file names are as follows: `results/AB/AB+bb-t18+fp-f17-r21+dd+rrf-d10000+alphadiversity.tsv` `results/AB/AB+bb-t18+fp-f17-r21+dd+cls-gg+rrf-d10000+beta_braycurtis.tsv` `results/AB/AB+bb-t18+fp-f17-r21+dd+cls-gg+rrf-d10000+beta_weightedunifrac.qza`
- **Summary:** Having in mind the need of users to link the analysis made on snaq to other software tools, we prepared a series of special targets that generate results ready to be used elsewhere:
  – Phyloseq: Generates a Phyloseq object in RDS file, that can be easily imported to an R environment for subsequent analysis steps. This object includes the ASV table, taxonomy, and phylogenetic tree without rarefaction.
  – Biom: Produces biom table with taxonomy after rarefaction.
  – Manta: produces manta ready input files that can be easily uploaded in Manta for results storage and further analysis.
  Finally, a special `zip` file can be produced, as the one shown in Figure 1, that includes all content summarized in Table 2. A complete list of the files produced for an example analysis process is provided in Supplementary Table (1).
- **Quality control:** An additional, optional step, that runs fastqc (Andrews 2010) and/or multiqc (Ewels et al. 2016). Unlike with other targets, the results of this stage are saved to a different `quality` folder, in our example: `result/AB/quality`. It is executed by using targets in the following form: `results/AB/quality/AB+bb-t18/multiqc/` `results/AB/quality/AB/multiqc/` Quality control can be applied either to the original data `AB`, or to any intermediate result obtained before the use of the DADA2 step. A special case that combines all fastqc reports in all subfolders of `quality` folder into a new `quality_summary` folder is achieved by using the target `results/AB/quality_summary/`.

**Table 2:**
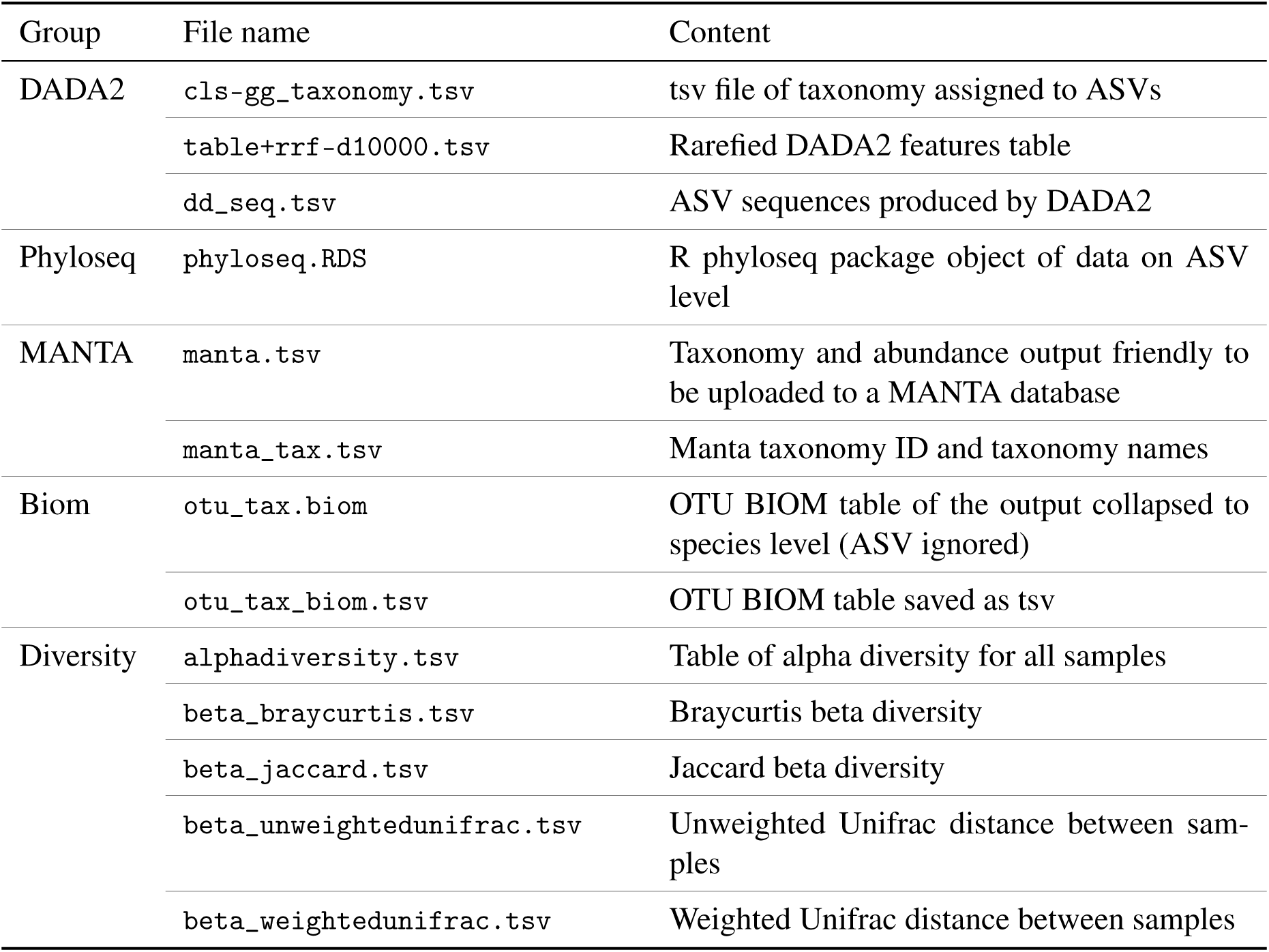
Summarized output, all the files are preceded by the parameters for the analysis.

### Installation

The only prerequisite on Linux or Mac is the installation of both Conda (*Anaconda Software Distribution* 2020) and Mamba (*Mamba, The fast cross platform package manager* 2021). We recommend running Snaq after activating the Snakemake environment installed using Conda. Conda and Mamba are also required to facilitate the installation of QIIME2 and other required tools automatically when Snaq is run for the first time. Docker installation is the only requirement in running Snaq on Windows using a docker container. Figure 3 shows the file structure after installation.

**Figure 3:**
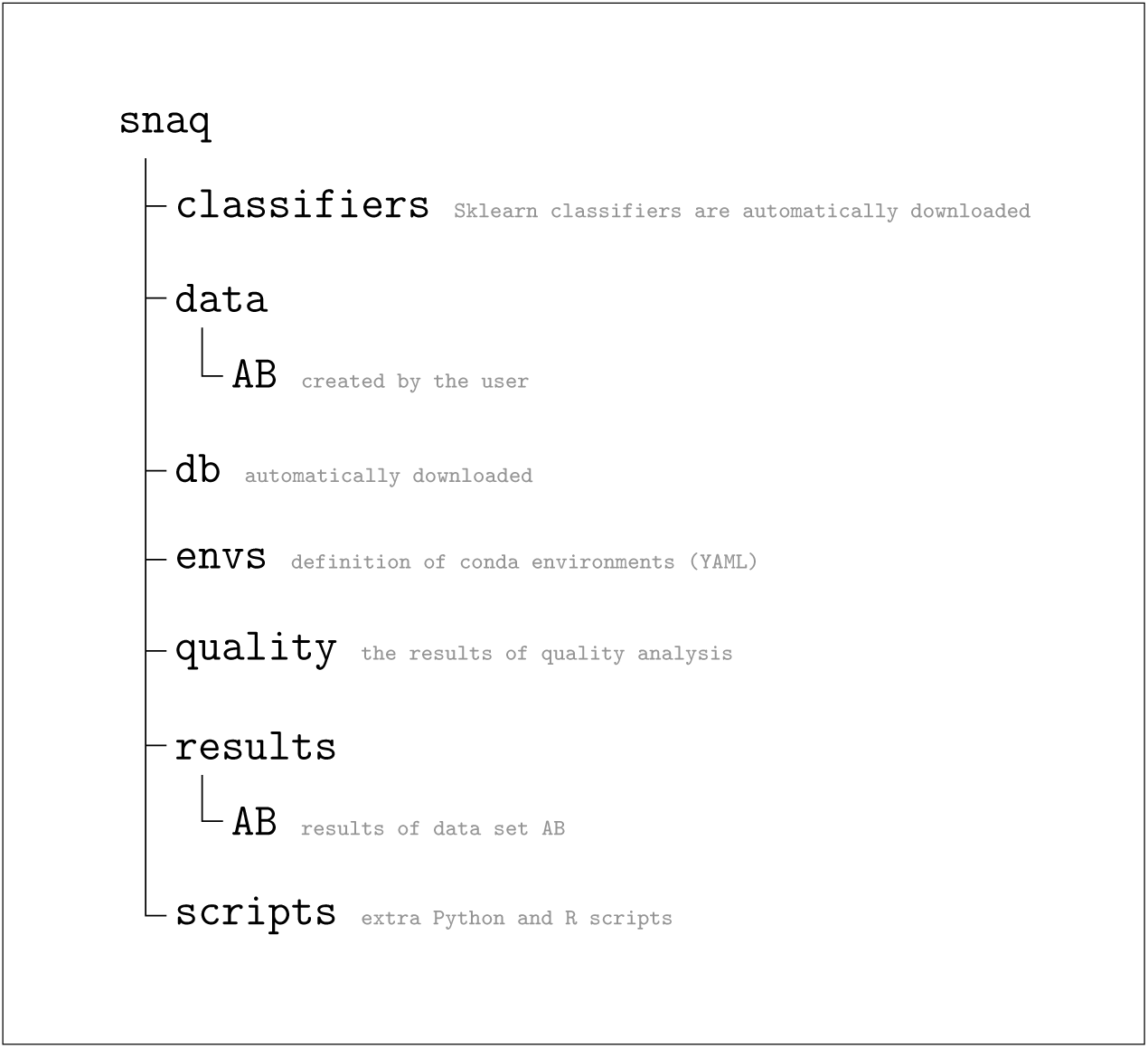
Folder structure of Snaq after installation. Contents of data and results folders will vary according to use after installation. Notice that all data sub-folders need to be named using capital letters.

The input data of Snaq are paired-end FASTQ files. Snaq automatically distinguishes pair ends by one of two identifiers `_R1_` or `_1.fastq`. If other identifiers are used, a manifest file needs to be prepared and saved as `results/AB/AB_manifest.tsv` following the QIIME2 manifest file instructions. If this file is present, the first step of creating a manifest file will be ignored.

## Results and Discussion

Snaq is designed to wrap QIIME2 processing of paired-end FASTQ files generated by Illumina sequencers to help automation, optimization, and take care of the data storage. It requires minimal effort in installation and configuration; moreover, it can run on all major operating systems. The user only needs a single command to run the pipeline defining required parameters for the analysis in the target file name.

Snaq can be installed directly from GitHub into a user specified location of choice. Notice that the installation directory needs to have enough free space to allocate for all input and intermediate data sets, together with all final results for any particular analysis. Free space is also required for the software programs and databases used in the analysis.

Input data must be saved inside the data folder `<snaq folder>/data/` after creating a new folder with the dataset name inside it. Dataset names should consist of capital letters without numbers or special characters in order to avoid confusing them with terms reserved to represent different pipeline stages. Once input data is available and the first step in the analysis is executed, Snaq (through Snakemake) will automatically build the Conda environment required for that step and download the QIIME2 plugins specified in the corresponding environment YAML file. Environment description files are located in the `<snaq folder>/envs/` folder. Notice that this makes the installation of necessary software and the download of taxonomy classifiers an automatic process, only to be performed the first time it is required.

Although Snaq does not cover all the possible uses of QIIME2 and related platforms in 16S data analysis, it provides a complete pipeline that can be extended by adding new rules or modifying the currently available ones. Moreover, following the descriptive target file name strategy makes it easier for Snaq to decide which step to run and skip. That also gives the developer who wants to modify Snaq the freedom to modify the pipeline and add new rules besides the current ones, as a different sequence of rules can be followed depending on the target file name. Other classifiers can also be added to the `classifiers/` folder, and the file name can be used in the target file name; if a classifier is named “abc” and saved as `abc-classifier.qza`, then we can use it with this target: `results/AB/AB+bb-t18+fp-f17-r21+dd+cls-abc+rrf-d1000.zip` without any modifications.

Other pipelines automate 16S data analysis. For instance, Fung et al. showed multiple protocols to run QIIME2 using QAP (QIIME2 automation pipeline) shell scripts. In addition, their paper gave detailed explanations of many steps and descriptions of their results. Multiple commands need to be executed to run the analysis using QAP; moreover, it provides more options and different approaches than Snaq follows (Fung et al. 2021). Estaki et al. provided a comprehensive description of the QIIME2, showing several steps of QIIME2 analysis starting from data import to final steps, including Jupiter notebooks (Estaki et al. 2020). Hu and Alexander implemented a Snakemake pipeline for QIIME2 analysis (Hu and Alexander 2020). It is designed to run with parameters predefined in the configuration file; hence changing the parameters requires the modification of manifest and configuration files. Dadasnake is a pipeline that automates DADA2 analysis outside the setting of the QIIME2 framework (Weißbecker et al. 2020).

Compared to the pipelines mentioned above, Snaq allows dynamic modification of key parameters by modifying the target file name. It also provides a more straightforward installation process and clear output. Moreover, Snaq allows running multiple data sets in the same pipeline setting by having multiple folders in the `data` folder.

The concept of a descriptive output file name allows high freedom for the pipeline extension. For example, suppose a new tool is added to the pipeline, such as trimmomatic. In that case, the user must add a new rule in Snakefile, following the snakemake approach, and give it a unique identifier. For example, let us say “tm” then add the key parameters that he is interested in modifying so that the target could become +tm-p12+; then, this part can be incorporated into the whole pipeline. An example target name is a `results/AB/AB+tm-p12+bb-t18+dd+cls-gg+rrf-d1000.zip`, and that will allow the user to have another option (trimmomatic) quickly incorporated. The user needs, of course, to be careful regarding making the output of that new rule usable in the following rule. We plan to improve this pipeline in the future by integrating other tools or producing the results in other file formats and illustrations. That, of course, will be driven by our research demands; however, we are also willing to help and welcome users who wish to add or contribute by adding new components to the pipeline.

## Conclusion

We have introduced Snaq, a Snakemake pipeline for QIIME2 16S data analysis, including data QC and trimming.

Snaq is designed to be dynamic by using a customly specified target file naming system. Modifying key parameters within the target name also helps the user efficiently perform a series of iterative analyses, taking automatic advantage of previously calculated intermediate steps and keeping track of results.

Installation and running of Snaq are easy. Moreover, Snaq can be extended according to the users’ needs by adding new rules.

## Author Contributions

Conceptualization: AM, KM; Methodology: AM, YC, KM; Software: AM; Validation: AM, CH, YC; Writing-Original Draft: AM; Writing-review and editing: AM, KM, YC, RA; All the authors reviewed the final draft and approved the submission.

## Acknowledgments

We are grateful to Jonguk Park, Hitoshi Kawashima, Lokesh P. Tripathi for their valuable suggestions and help.

## Supplementary Material

Supplementary table 1.

## Data Availability Statement

The code is available in GitHub: https://github.com/attayeb/Snaq, along with installation and usage instructions.

## Footnotes

1 For details on the architecture of QIIME2 visit https://dev.qiime2.org/latest/architecture/. Last accessed: January 24th, 2022.

2 If a rule named “all” is defined, it is possible to run a predefined workflow without the specification of a target file.

3 Some tools required by QIIME2 do not run natively on Windows environments. Windows users could potentially use snaq through containers, windows Linux subsystem, and/or virtual machines.

4 Even when the command indicates the location of the result files, this also automatically maps to input folder under data/ with the given name

5 All examples assume the use of an input dataset, located in a source folder named data/AB folder

