## Supplemental Table 1 for "Snaq: A Dynamic Snakemake Pipeline for Microbiome data analysis with QIIME2"

| stage | file name |
| --- | --- |
| import data | AB_manifest.tsv<br>AB.qza |
| quality trimming | AB+bb-t18.qza |
| primer cropping | AB+bb-t18+fp-f17-r21.qza |
| | AB+bb-t18+fp-f17-r21+dd_seq.qza<br>AB+bb-t18+fp-f17-r21+dd_seq.tsv<br>AB+bb-t18+fp-f17-r21+dd_stats.qza<br>AB+bb-t18+fp-f17-r21+dd_table.qza |
| taxonomy assignment | AB+bb-t18+fp-f17-r21+dd+cls-gg_taxonomy.qza<br>AB+bb-t18+fp-f17-r21+dd+cls-gg_taxonomy.tsv<br>AB+bb-t18+fp-f17-r21+dd+cls-gg_asv.biom |
| phylogeny tree | AB+bb-t18+fp-f17-r21+dd+fasttree.nwk<br>AB+bb-t18+fp-f17-r21+dd+fasttree_rooted.qza<br>tree/ |
| phyloseq | AB+bb-t18+fp-f17-r21+dd+cls-gg+phyloseq.RDS |
| rarefaction | AB+bb-t18+fp-f17-r21+dd_table+rrf-d10000.qza<br>AB+bb-t18+fp-f17-r21+dd_table+rrf-d10000.tsv |
| diversitry | AB+bb-t18+fp-f17-r21+dd+rrf-d10000+alphadiversity.tsv<br>AB+bb-t18+fp-f17-r21+dd+cls-gg+rrf-d10000+beta_braycurtis.tsv<br>AB+bb-t18+fp-f17-r21+dd+cls-gg+rrf-d10000+beta_jaccard.tsv<br>AB+bb-t18+fp-f17-r21+dd+rrf-d10000+beta_unweightedunifrac.qza<br>AB+bb-t18+fp-f17-r21+dd+rrf-d10000+beta_unweightedunifrac.tsv<br>AB+bb-t18+fp-f17-r21+dd+rrf-d10000+beta_weightedunifrac.qza<br>AB+bb-t18+fp-f17-r21+dd+rrf-d10000+beta_weightedunifrac.tsv |
| manta | AB+bb-t18+fp-f17-r21+dd+cls-gg+rrf-d10000+manta_tax.tsv<br>AB+bb-t18+fp-f17-r21+dd+cls-gg+rrf-d10000+manta.tsv |
| biom | AB+bb-t18+fp-f17-r21+dd+cls-gg+rrf-d10000+otu_tax.biom<br>AB+bb-t18+fp-f17-r21+dd+cls-gg+rrf-d10000+otu_tax_biom.tsv<br>AB+bb-t18+fp-f17-r21+dd+cls-gg+rrf-d10000+otu_tax.qza |
| summary | AB+bb-t18+fp-f17-r21+dd+cls-gg+rrf-d10000.zip |

**Supplementary Table 1:** a complete list of all produced files for an imaginary data set "AB".
